# Light-Activated Nanoscale Gas Vesicles Selectively Kill Tumor Cells

**DOI:** 10.1101/771881

**Authors:** Ann Fernando, Jean Gariépy

## Abstract

Protein-based nanobubbles such as halophilic archaeabacterial gas vesicles (GVs) represent a new class of stable, homogeneous nanoparticles with acoustic properties that allow them to be visualized by ultrasound (US) waves. To design GVs as theranostic agents, we modified them to respond to light, with a view to locally generate reactive oxygen species that can kill cancer cells. Specifically, up to 60,000 photoreactive chlorin *e*6 (C*e*6) molecules were chemically attached to lysine ε-amino groups present on the surface of each purified *Halobacterium sp.* NRC-1 GV. The resulting fluorescent NRC-1 C*e*6-GVs have dimensions comparable to that of native GVs and were efficiently taken up by human breast [MCF-7] and human hypopharyngeal [FaDu-GFP] cancer cells as monitored by confocal microscopy and flow cytometry. When exposed to light, internalized C*e*6-GVs were 200-fold more effective on a molar basis than free C*e*6 at killing cells. These results demonstrate the potential of C*e*6-GVs as novel and promising nanomaterials for image-guided photodynamic therapy.

## Introduction

Image-guided therapies are precision medicine-based treatments being developed in many cases to treat cancer patients. These therapies benefit from the development of theranostic agents that integrate both targeted diagnostic and therapeutic functions.^1,2,3^

Theranostics often incorporate radionuclides or MR contrast agents as imaging probes in conjunction with cytotoxic nuclides or drug-conjugates linked to a tumor-targeting agent such as a small ligand or an antibody that delivers the payload. An alternate design strategy to reduce potential off-target toxicities arising from delivering these agents systemically is to create theranostics that are only activated within the tumor environment (pro-drug like theranostics that become activated as a result of a local change in pH or by tumor-associated proteases) or upon exposing tumors to an external, focused energy source that would remotely activate such agents within the tumor environment only. Nanoparticles are particularly suited to this purpose as they tend to accumulate within the abnormal tumor neovasculature as a consequence of the enhanced permeability and retention (EPR) effect. ^4,5^ Tumor selectivity can be encoded into nanoparticles by making them sensitive to an externally applied, ultrasound and light source; physical methods that are presently being used in image-guided therapy. Protein-based gas vesicles (GVs) represent an example of a naturally occurring nanostructure that could be used for this purpose. In the present study, we focused on GV nanostructures genetically encoded by the aquatic halophilic archeon *Halobacterium salinarum* NRC-1.^6^ They are composed of two dominant subunits termed gas vesicle protein A (GvpA) and gas vesicle protein C (GvpC). ^6^ These GVs have recently been shown to respond to ultrasound waves and to produce hyperpolarized magnetic resonance contrast for imaging.^7,8^ In the present study, we explored the potential of such GVs as photodynamic therapy (PDT) agents. We report the design of a light-activated GV nanoparticle where the photo-reactive dye chlorin *e*6 (C*e*6) was coupled to free amino groups on the surface of GVs. The resulting nanoscale size structures were found to readily accumulate into cancer cells and kill them in response to light activation displaying remarkable enhancements in cytotoxicity towards cancer cells relative to free C*e*6.

## Results and Discussion

### Preparation and characterization of GVs modified with chlorin *e*6 (C*e*6-GVs)

Gas vesicles (GVs) are air-filled nanobubbles that have been shown to respond to ultrasound (US) waves.^7^ In this study, GVs were isolated from the *Halobacterium sp.* NRC-1 to serve as nanocarriers for the light-activated drug, chlorin *e*6 (C*e*6) with a view of creating therapeutic GVs beyond their US-imaging properties. As such, NRC-1 GVs were purified and their surface modified with C*e*6 to demonstrate their ability to be internalized by cancer cells and to enhance the cytotoxicity of C*e*6 as a photodynamic therapy (PDT) agent. C*e*6-modified GVs represent a first attempt at generating GV-based theranostic agents aimed at killing cancer cells. Specifically, C*e*6 absorbs light with maxima observed at 400 nm and 660 nm. The photodynamic effect occurs upon absorption in the red region of the spectrum where human tissue is optically transparent, reaching tissue depth of up to 16 mm.^9,10^ C*e*6 is also a fluorescent compound, which enables the use of optical imaging to detect its accumulation into tumors.^11^

*Halobacterium sp.* NRC-1 GVs used in this study are composed of two major protein species (GvpA and GvpC) as well as several minor proteins involved in the nanobubble assembly. Importantly, GvpA accounts for >90% of the nanobubble structure and shell^6^. GvpC binds non-covalently to the surface of the GV shell. However, it is stripped from the nanobubble surface during the process of isolating GVs from the halobacterium, as confirmed by mass spectrometry of our GV preparations (Supplementary Figure S1). As such, the two lysine ε-amino groups present in GvpA (lysine 19 [helix I] and lysine 60 [helix II] of GvpA; depicted in Fig. 1A) represent the main sites for covalently attaching C*e*6 to the surface of GVs.

**Fig. 1.**
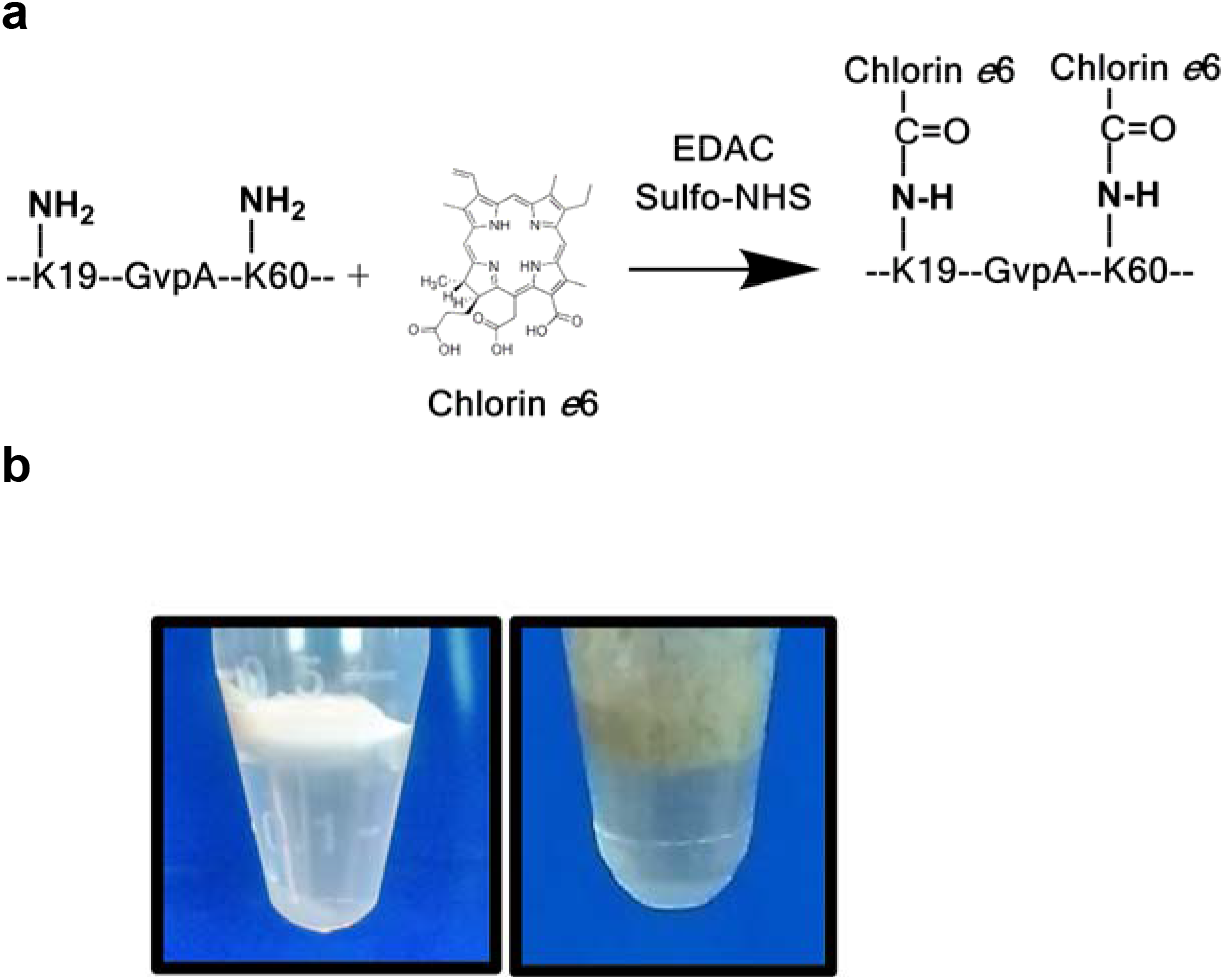
A) Diagram depicting the conjugation of chlorin *e*6 (C*e*6) carboxylic groups to the two lysine ◻-amino groups (K19 and K59) of GvpA, the main protein component of *Halobacterium sp. NRC-1* Gas Vesicles (GVs). B) GVs are air-filled protein nanobubbles recovered by flotation from lysed halophilic archaeabacteria. Purified floating layers are showed of wild type (WT) *Halobacterium sp. NRC-1* GVs (white) and C*e*6-modified GVs (green-grey color). ***(Image Fitting = 1 column)***

Experimentally, wild type gas vesicles (WT GV) from the *Halobacterium sp.* NRC-1 were isolated by flotation from lysed halobacteria and purified by repeated centrifugation steps as described elsewhere (Fig. 1B). ^12^ The carboxylic groups of chlorin *e*6 were then activated with EDAC and sulfo-NHS to form esters that reacted with the two free ε-amino groups of GvpA on WT GVs, resulting in an olive-green colored nanobubble preparation termed C*e*6-GVs (Fig. 1B).

The particle size of these GVs was studied by dynamic light scattering (DLS) and transmission electron microscopy (TEM). Both WT GVs and C*e*6-GVs adopted a characteristic prolate-spheroid, lemon-shaped structure of comparable dimensions as confirmed from electron micrographs, suggesting that decorating the surface of GVs with Ce6 molecules minimally affected particle structure and size (Fig. 2, Table 1). The length of C*e*6-GVs along their long and short axes were measured to be 417 nm and 196 nm respectively according to TEM (Fig. 2, Table 1). Estimates of WT GVs and Ce6-GVs particle diameter by DLS (273±3 nm and 338±39 nm respectively; Table 1) assumes that GVs are spherical. The calculated diameters approximate the average length of both their short and long axes as determined by TEM (Table 1). The zeta potential of C*e*6-GVs was significantly lower (−30 mV) than WT GVs (−2.4 mV) as C*e*6 contains three negatively charged carboxylic acids (Table 1). The increased negative charge suggests that C*e*6-GVs may have good colloidal stability due to repulsion between the nanoparticles.

**Fig. 2.**
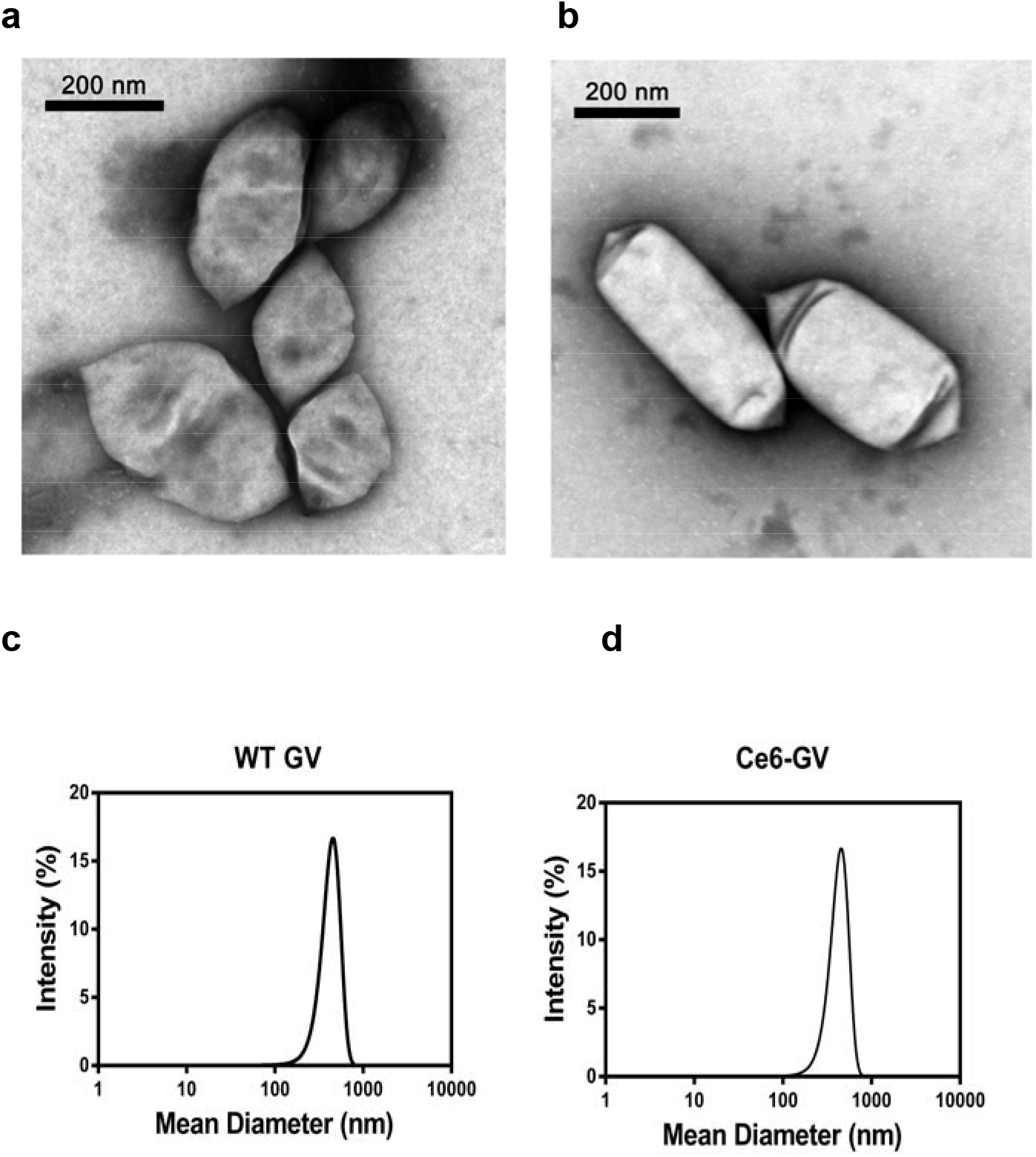
Characterization of WT GVs and C*e*6-labeled GVs. Transmission electron micrographs of A) WT GVs and B) C*e*6-labeled GVs indicating that modifying their surface with C*e*6 yields comparable nanostructures. Similar dynamic Light Scattering size distribution profiles are observed for C) WT GVs and D) C*e*6-GVs. ***(Image Fitting = 1.5 column)***

**Table 1.**
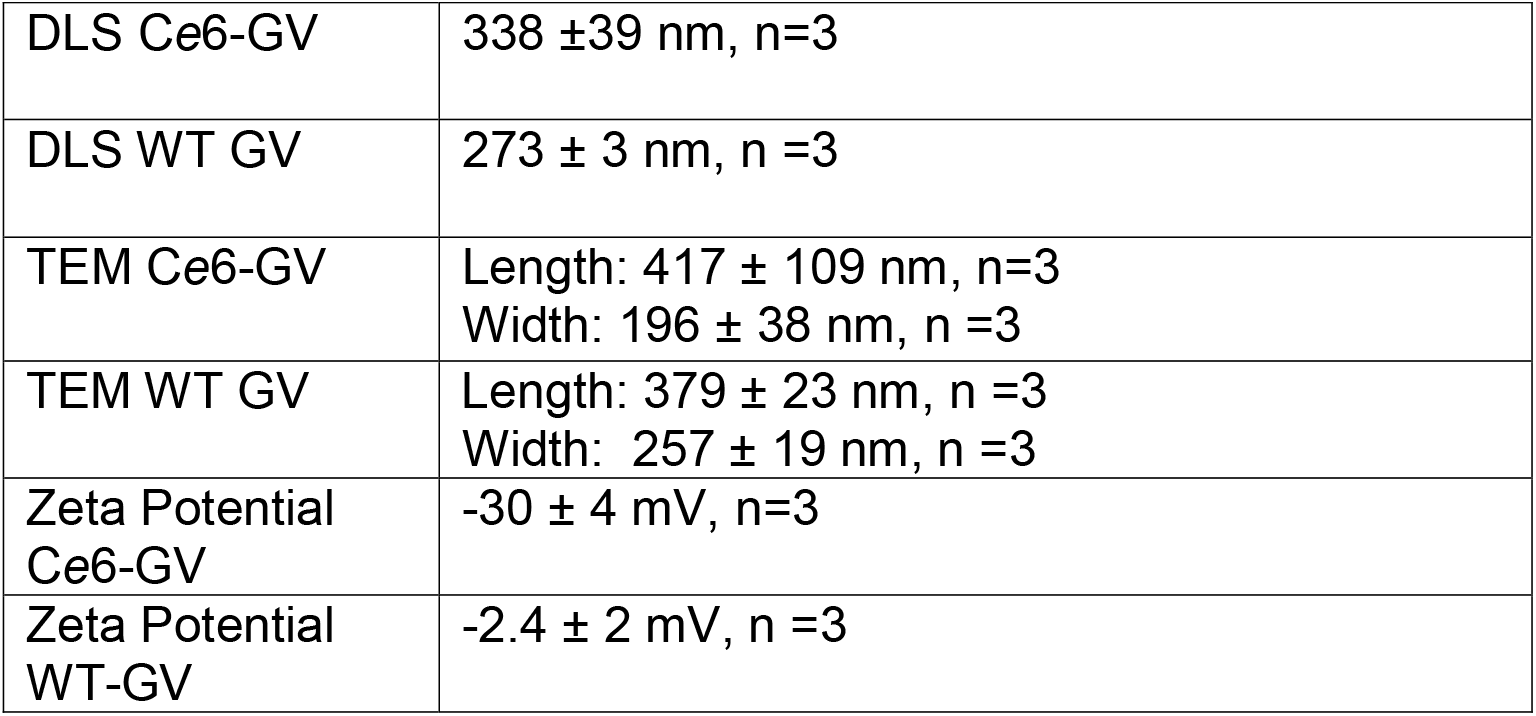
Summary of size parameters for WT GVs and C*e*6-GV. Results are shown as mean ± S.D.***(Image Fitting = 1 column)***

|  |  |
| --- | --- |
| DLS Ce6-GV | 338 ± 39 nm, n=3 |
| DLS WT GV | 273 ± 3 nm, n =3 |
| TEM Ce6-GV | Length: 417 ± 109 nm, n=3<br>Width: 196 ± 38 nm, n =3 |
| TEM WT GV | Length: 379 ± 23 nm, n =3<br>Width: 257 ± 19 nm, n =3 |
| Zeta Potential Ce6-GV | -30 ± 4 mV, n=3 |
| Zeta Potential WT-GV | -2.4 ± 2 mV, n =3 |

Using surface area measurements of WT GVs based on TEM micrographs (Fig. 2 A,B; Table 1; ~380 nm by 260 nm) along with the experimentally determined surface area of one GvpA molecule, the main repeating structural component of the nanobubble shell, ^13^ the mass of one *Halobacterium* GV was estimated to be 444 MDa (See Supplementary Figure S2) with a single NRC-1 GV being composed of ~55,000 GvpA proteins. Since the two lysines present in GvpA (K19 in helix I and K60 in helix II; Fig. 1A) are projected to be solvent-accessible based on solid-state NMR,^14^ one can project that up to 110,000 ε-amino groups are available to react with an amine reactive form of C*e*6. To accurately determine the extent of C*e*6 conjugated to WT GVs, we measured the total amount of protein (amino acid analysis) and released C*e*6 content by hydrolyzing Ce6-GVs samples in 6N HCl. It was estimated from hydrolysates that ~60,000 molecules of C*e*6 molecules were coupled per nanobubble; a value representing a loading efficiency of approximately one C*e*6 molecule per GypA protein.

The presence of C*e*6 on GVs was further confirmed by recording the fluorescence emission spectra of C*e*6 released by acid hydrolysis from C*e*6-GVs (Fig.3). The observed fluorescence emission maximum at 660 nm recorded for C*e*6-GVs was comparable to that of free C*e*6, while hydrolysates from WT GVs were non-fluorescent (Fig. 3). This finding is comparable to the published fluorescence emission spectra of equimolar doses of C*e*6-liposomes and free C*e*6^15^ or C*e*6-albumin nanoparticle formulations.^16^ These results indicate that C*e*6 can be efficiently loaded onto GVs and also demonstrate that conjugation to GVs does not significantly alter the spectral properties of C*e*6, nor the shape and dimensions of GVs.

**Fig. 3.**
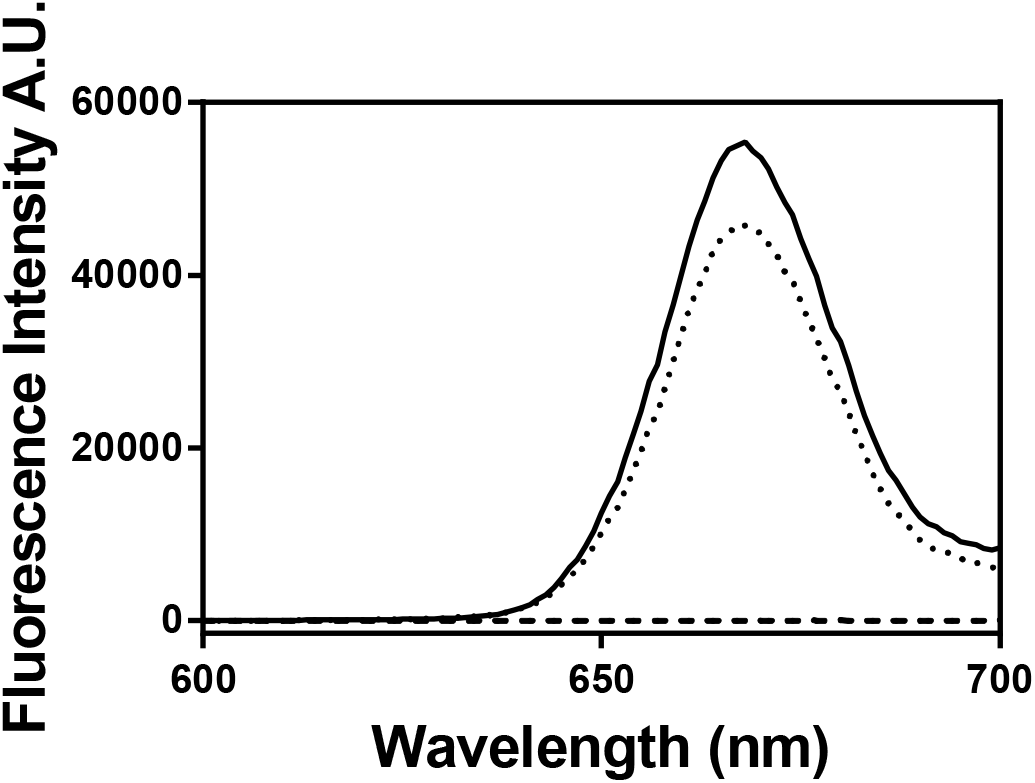
The fluorescence spectrum for C*e*6 released from acid-hydrolyzed C*e*6-GVs was comparable to that recorded for free C*e*6. The emission spectra were recorded between 600 and 700 nm (◻_exc_ 400 nm) at 1 nm interval. The spectrum recorded for WT GVs hydrolysates confirmed the absence of a C*e*6 spectral signature. All spectra are representative of recordings from two independent acid-hydrolyzed samples. ***(Image Fitting = 1 column)***

### C*e*6-GVs are taken up by cancer cells and accumulate in endosomes

The light-induced cytotoxicity of C*e*6-containing GVs depends on their ability to be internalized by cancer cells. Upon reaching the cytoplasm and following light activation, C*e*6 molecules are excited to their singlet→triplet states where their energy can be transferred to O_2_ to generate reactive singlet oxygen (^1^O_2_).^17^ Alternatively, activated C*e*6 can react directly with proteins, nucleic acids and lipids to form reactive oxygen species (ROS; superoxide anion, hydroxyl radical, hydrogen peroxide) causing oxidative damage leading to cell death.^17,18^

The cellular uptake of C*e*6-GVs, WT GVs and free C*e*6 by human breast carcinoma MCF-7 and human hypopharyngeal squamous cell carcinoma FaDu cells was thus assessed by flow cytometry and confocal microscopy. For both techniques, fluorescence emission signals arising from the chlorin *e*6 chromophore (λ_exc_ 403 nm) were captured between 660-680 nm (Fig. 3). The intracellular uptake of C*e*6-GVs and free C*e*6 was monitored by flow cytometry in terms of mean fluorescence intensities (MFI) as a function of time and temperature (Fig. 4). As expected, cellular uptake did not take place at 4 °C for either cell lines (Fig. 4 A,C) while the internalization of C*e*6-GVs at 37 °C reached a plateau at 8 hours and 22 hours for the FaDu-GFP and MCF-7 cell lines respectively (Fig.4 B,D). In contrast, free C*e*6 minimally enters these cells while no fluorescence signal could be detected for WT GVs (Fig. 4). These results show that conjugation of C*e*6 to GVs improves the intracellular delivery of this drug into cancer cells. Cellular uptake of other C*e*6 nanoparticles have been previously described as exemplified by C*e*6-octalysine conjugated to superparamagnetic iron oxide nanoparticles being taken up by SKOV3 cells relative to free C*e*6.^19^

**Fig. 4.**
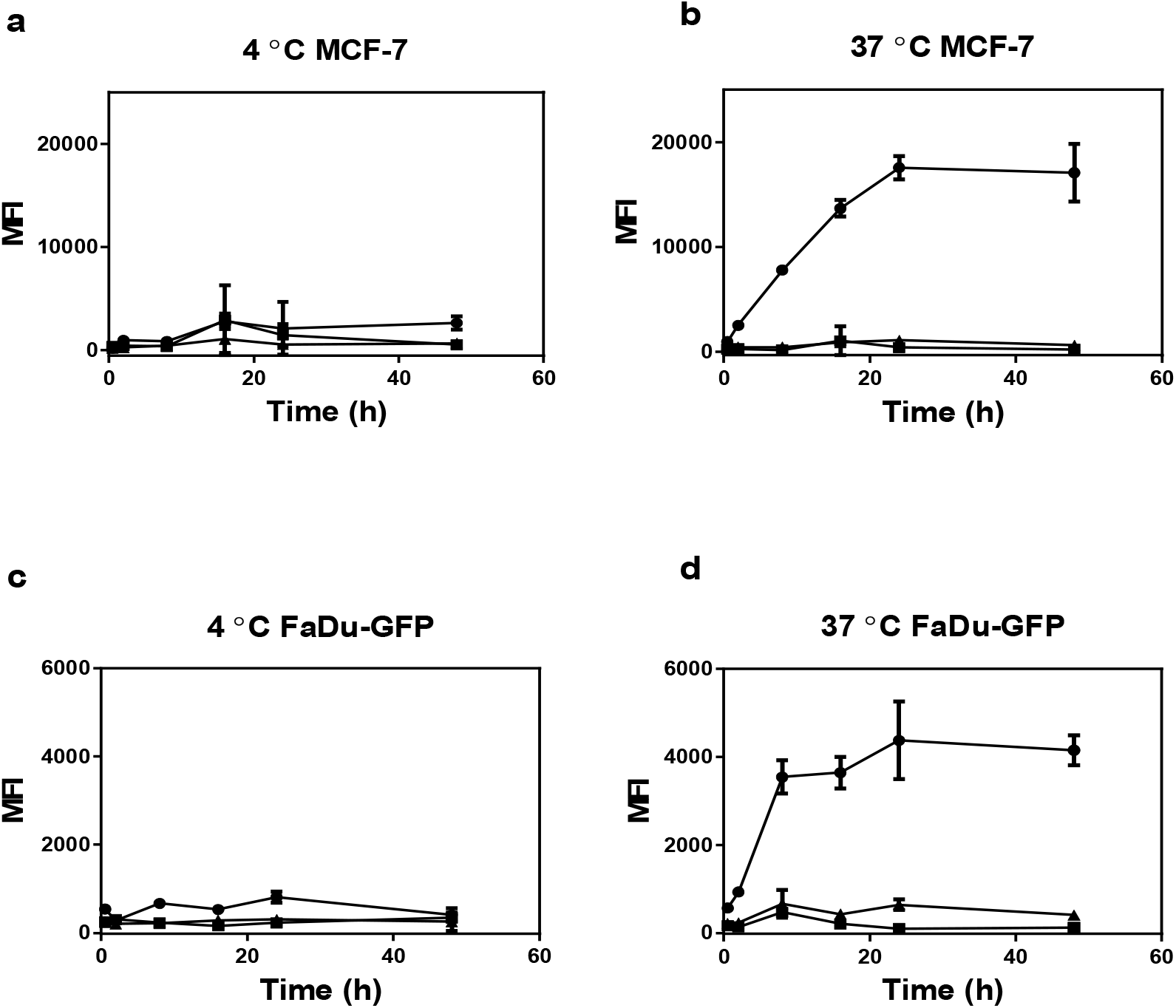
Mean fluorescence intensities (MFI) as determined by fluorescence-activated cell sorting (FACS) indicate that C*e*6-GVs are internalized as a function of time by MCF-7 cells and FaDu-GFP cells at 37 °C (Panels B and D). In contrast, minimal cellular uptake takes place at 4°C (Panels A and C). Cells were equivalently treated with 670 nM of C*e*6 either as C*e*6-GVs (closed circles) or as the free drug (closed triangles) over a 48-hour period. As projected, equivalent concentrations of WT GVs (closed squares) relative to C*e*6-GVs did not generate significant MFI values. Results shown are representative of two independent trials performed in triplicate and presented as averaged MFI ± S.D. ◻_exc_ 405 nm; emission Filter 660-662 nm. ***(Image Fitting = 2 column)***

The cellular uptake of C*e*6-GVs at 37°C into MCF-7 and FaDu-GFP cells was further confirmed by confocal microscopy (Fig. 5). Specifically, a punctate pattern of C*e*6 fluorescence (in red) is observed in the cytosol of these cells after an 8-hour incubation period suggesting their compartmentalization into organelles such as endosomes or vacuoles. C*e*6-GVs did not reach the cell nucleus (Hoescht dye; blue color nuclei; Fig. 5 A,E). This cytoplasmic distribution pattern has been observed for other C*e*6-conjugates and nanoparticles.^9, 20^

**Fig. 5.**
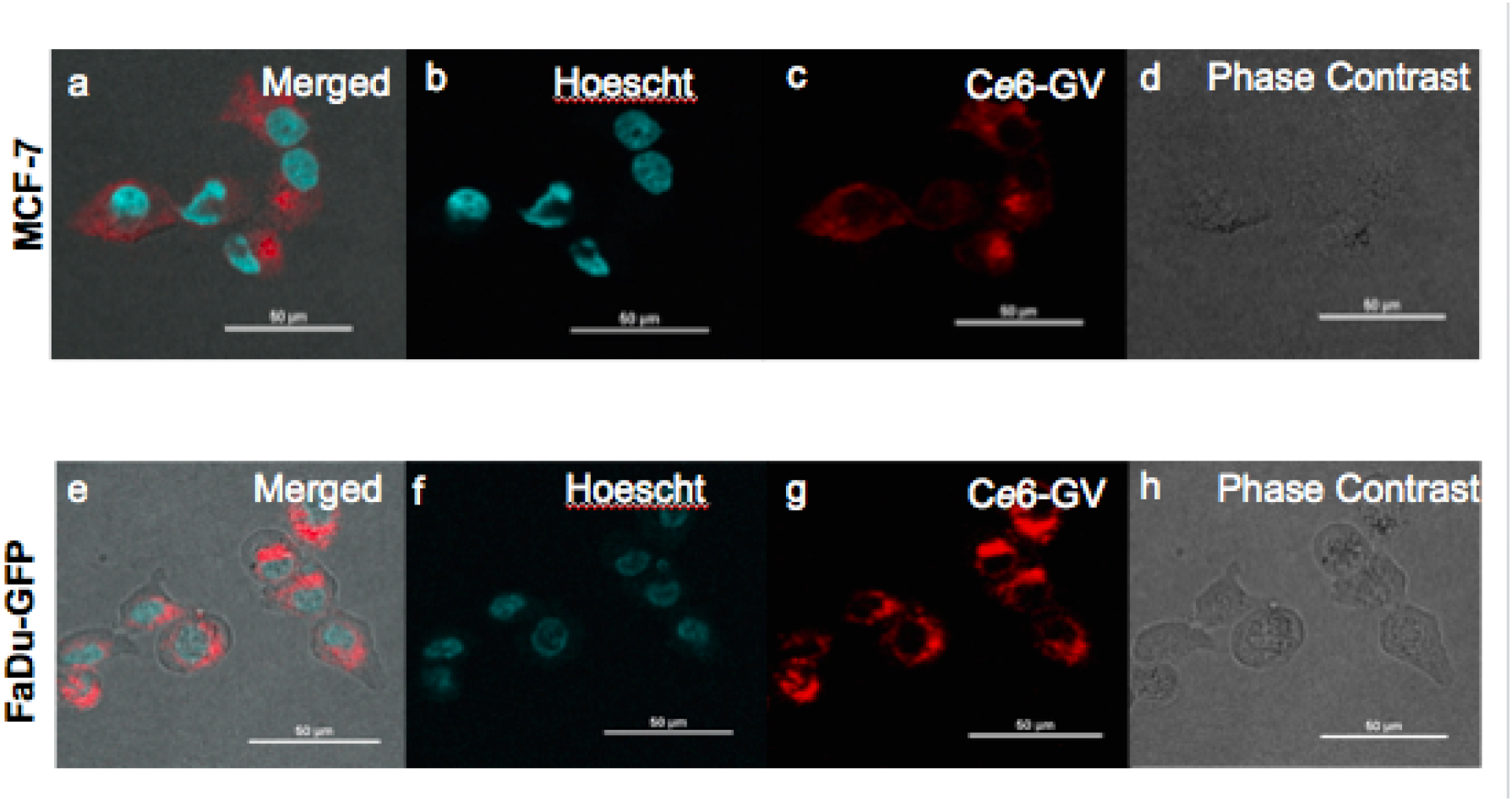
Merged confocal fluorescence and phase contrast microscopy images indicate that C*e*6-GVs accumulate in the cytoplasm of cancer cells in a punctate- like pattern (in red) suggesting their entrapment in endocytic vesicles or vacuoles. The lack of co-localization of C*e*6-GVs with the cell nucleus (Hoescht dye; blue stain) suggests that reactive oxygen species generated by C*e*6-GVs being exposed to light would mainly accumulate in the cytoplasmic compartment of MCF-7 (panels A-D) or FaDu-GFP (panels E-H) cells after 8 hours of incubation. ***(Image Fitting = 1.5 column)***

### Gas Vesicles dramatically enhance the phototoxicity of C*e*6 towards cancer cells

Human MCF-7 breast cancer and FaDu-GFP pharyngeal cancer cells were exposed to increasing concentrations of C*e*6-GVs, free C*e*6, or an equivalent amount of WT GVs for 24 hours, followed by a 10-minute exposure to light. Their viability was assessed 24-hours later and reported as a function of the molar concentration of C*e*6 being given (Fig. 6). Both cell lines remain viable in the absence of light treatment except for cells exposed to high doses of C*e*6-GVs (10^−5^ M range; Fig 6 A,C). Upon light activation, cell viability was lost in a dose-dependent manner with C*e*6-GV being more potent than the free drug (Fig.6, B,D). Specifically, CD_50_ values towards both cell lines were determined for C*e*6-GV and free C*e*6 (Table 2) following light exposure, with the drug covalently attached to GVs being 200-fold more toxic on a drug molar basis than free C*e*6 (Fig. 6 B,D). WT GVs were only toxic towards MCF-7 or FaDu-GFP at very high concentrations (at concentrations >10^−4^ M relative to comparable C*e*6-GV doses given; Supplementary Figure S3) The enhanced toxicity observed for C*e*6-GV correlates with its greater cellular uptake seen in both cell lines in contrast to free C*e*6 (Fig. 4,5). This enhancement in light-activated toxicity relative to the free form of the drug has been observed in SKOV3 and MDA-MB-231 cells for other C*e*6-nanoparticles such as C*e*6-SPION ^19^ or C*e*6-conjugated poly(ethylene glycol)-poly-(d,l-lactide) ^21^ nanoparticles, likely owing to their improved cellular uptake. In contrast, the photosensitizing efficacy and toxicity of a C*e*6 conjugated human serum albumin nanoparticle was comparable to that of free C*e*6 ^22^ suggesting that all C*e*6-nanoparticle formulations are not equivalent in terms of enhanced cytotoxicity. Interestingly, agents such as C*e*6 can also be activated by ultrasound waves; an approach termed sonodynamic therapy that may allow one to target deep-seated tumors following the activation of the photosensitizer agent.^23^ For example, the viability of H22 hepatocellular carcinoma cells exposed to 50 μg/ml of free C*e*6 for 4 hours was decreased by 40 % following ultrasound treatment. ^24^ Although ultrasound is a less potent activation modality than light, our present study now suggests that the use of C*e*6-GVs rather than free C*e*6 may address this limitation. As such, C*e*6-GVs are now being assessed as a new class of sono-sensitive agents.

**Fig. 6.**
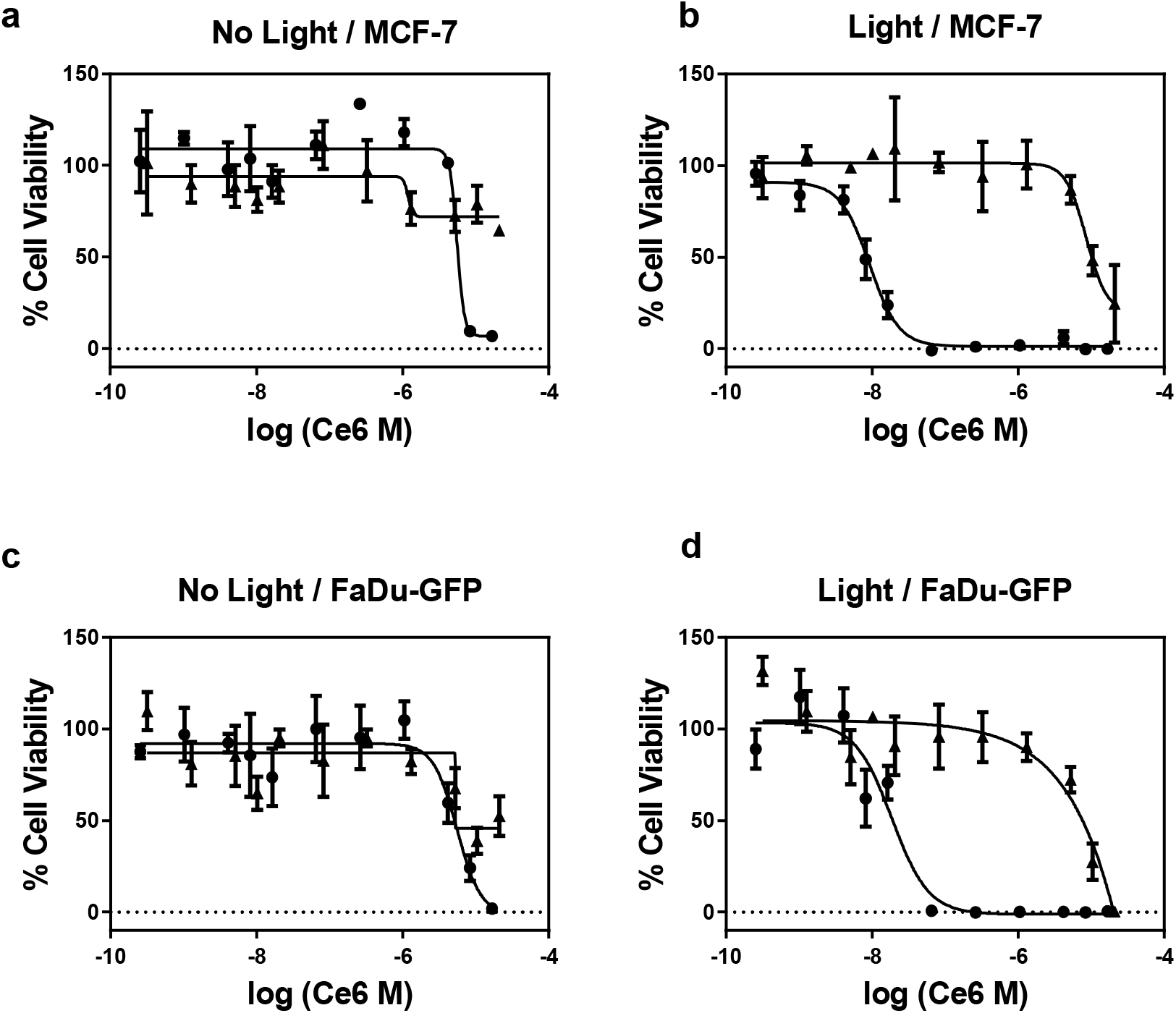
Dose-dependent cell viability study of MCF-7 cells or FaDu-GFP cells treated with free C*e*6 (closed triangles) or C*e*6-GVs (closed circles) and exposed (Panels B and D) or not (Panels A and C) to light as determined using the WST-1 cell viability assay. Results are shown as mean ± S.D. n=3 from 2 independent experiments, with each concentration performed in triplicate. ***(Image Fitting = 2 column)***

**Table 2.** Comparison of CD_50_ values determined for MCF-7 cells and FaDu-GFP cell lines following their exposure to free Ce6 and Ce6-GV. Molarity was calculated based on the molar concentration of C*e*6. ***(Image Fitting = 1 column)***

|  | MCF-7 CD<sub>50</sub> (M) | FaDu-GFP CD<sub>50</sub> (M) |
| --- | --- | --- |
| Ce6-GV | 2.6 ± 1.4 $\times 10^{-8}$ | 2.4 ± 0.8 $\times 10^{-8}$ |
| Free Ce6 | 5.3 ± 3.3 $\times 10^{-6}$ | 5.5 ± 4.0 $\times 10^{-6}$ |
| Free Ce6 CD<sub>50</sub> / Ce6-GV CD<sub>50</sub> | 204 | 229 |

### The enhanced toxicity of C*e*6-GVs towards cells is mechanistically related to their cellular accumulation and the production of intracellular ROS

The phototoxic effect of C*e*6 is related to the production of intracellular reactive oxygen species (ROS) as a consequence of the drug being exposed to light. This effect was measured by treating cells with the green fluorescence-emitting probe, DCFH-DA, prior to illumination. As presented in Fig. 7, the DCFH-DA fluorescence emission signal for MCF-7 and FaDu-GFP treated with C*e*6-GV confirm the presence of ROS in light-treated cells only (Fig 7A,B). For MCF-7 and FaDu-GFP cell lines, significantly more ROS were produced when C*e*6-GV-loaded cells were exposed to light as compared to C*e*6-GV-loaded cells kept in the dark (p = 0.0052 and p =0.0004 respectively). The difference in ROS production due to C*e*6-GV treatment was greater than ROS production due to free C*e*6 treatment in either MCF-7 (p= 0.0385) or FaDu-GFP cell lines (p= 0.0126). These results confirm that conjugating C*e*6 to GVs does not impair their capacity to produce ROS and is consistent with the superior uptake of C*e*6-GV relative to free C*e*6 demonstrated by flow cytometry (Fig. 4). Internalization events favoring the accumulation of C*e*6-GVs over the free drug inside cells probably play a dominant role in enhancing the toxicity of C*e*6 towards cancer cells. The generation of ROS is expected to cause damage to intracellular components such as protein, DNA, or membranes and trigger apoptotic or necrotic pathways to mediate cell death. ^25^ In the case of C*e*6-GVs, confocal images presented in Fig.5 suggest that these nanobubbles never reach the cell nucleus indicating that the generated reactive oxygen species following illumination are mainly deposited in the cytoplasm of these cells, preferentially causing the oxidation of lipids and proteins rather than damaging nucleic acid species. ^26, 27^

**Fig. 7.**
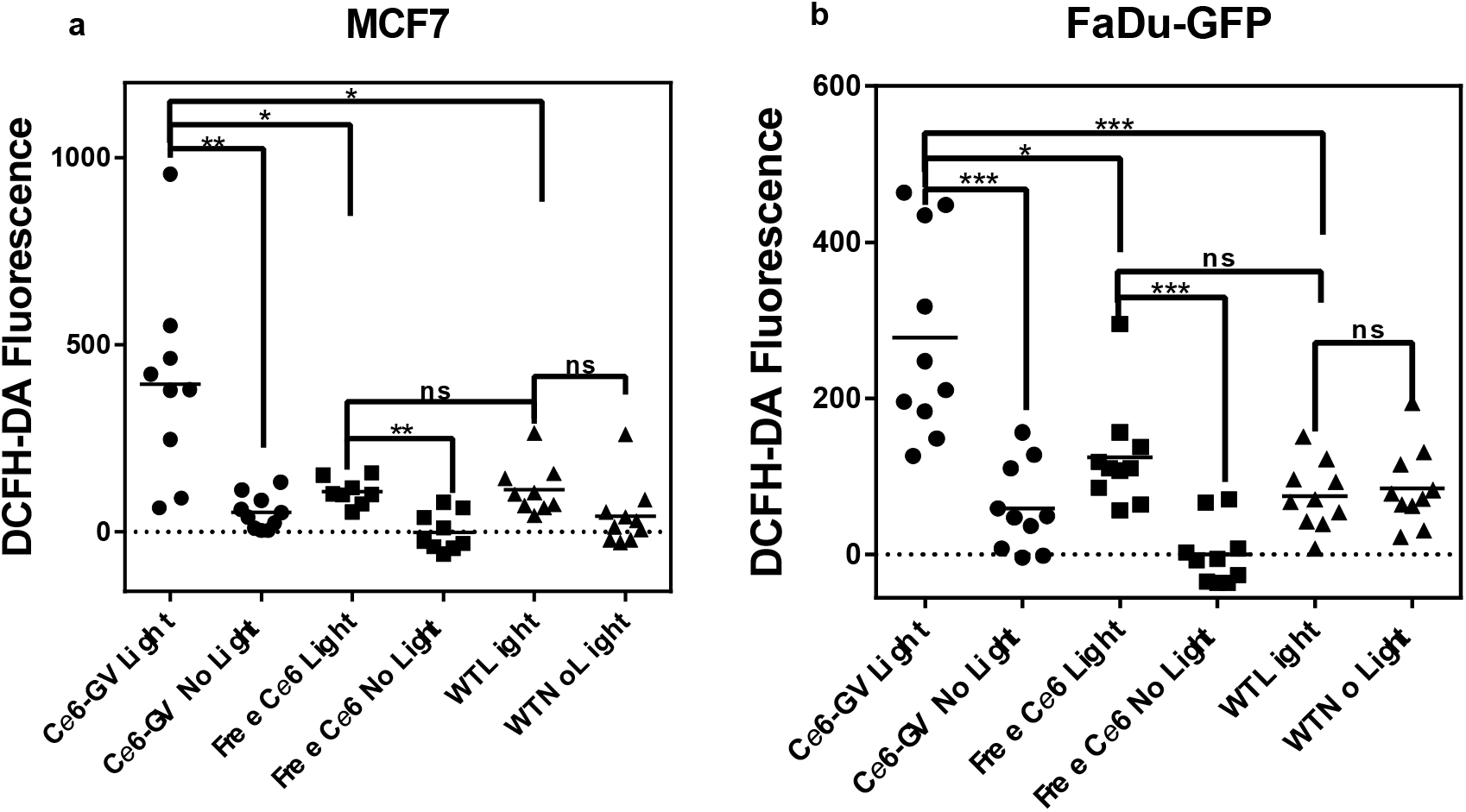
Production of Reactive Oxygen Species (ROS) in MCF-7 and FaDu-GFP cells treated with C*e*6-GV, WT GV, or free C*e*6. The ROS-sensitive probe DCFH-DA was subsequently loaded into treated cells with or without light activation of C*e*6. ROS levels were quantified using a fluorescence plate reader. The results are shown as mean ± S.D. (n=10). Statistical significance was determined with a paired t-test, p < 0.05*, p < 0.01**, p < 0.001 ***, p < 0.0001 ****. ***(Image Fitting = 1.5 column)***

## Conclusions

We have covalently linked the photoreactive drug C*e*6 to nanoscale gas vesicles (GVs) with a view to expand their potential as nanoparticles responding to ultrasound waves and now to light. This newly-introduced function suggests that GVs represent a good platform for designing stable theranostic agents that can be externally activated to serve as both imaging and now as locally-activated therapeutic agents. Specifically, C*e*6-GVs were efficiently internalized by MCF-7 and FaDu-GFP cancer cells and were highly effective in killing them *in vitro* upon light activation, relative to the free drug.

## Material and Methods

### Production and Purification of Gas Vesicles

Gas vesicles were isolated from *Halobacterium sp. NRC-1*. Cells were cultured in CM+ growth medium [4.3M Sodium Chloride, 81mM Magnesium sulfate heptahydrate, 10 mM trisodium citrate dehydrate, 27 mM potassium chloride, 0.5% casein hydrolysate (Sigma), 0.3% yeast extract (Difco)] at 42 °C, shaking at 100 rpm for one week until cells are confluent. The cultures were subsequently transferred to sterile separatory funnels and the fraction of cells expressing GVs at a high level were allowed to accumulate at the top by flotation over a one-week period. GVs were collected by hypotonic lysis of the buoyant cell fraction using 1 mM MgSO_4_ solution. After lysis, GVs were purified using repeated centrifugation steps at 300g overnight for at least three days and washed with PBS at each step. Intact buoyant vesicles were re-suspended in PBS and dialyzed against PBS (100 kDa MWCO dialysis membrane; Biotech CE) as a final step.

### Fabrication of chlorin *e*6-decorated GVs

Chlorin *e*6 (C*e*6; 119 μg, Cayman Chemicals, 20mM), 1-1-(3-di-methylaminopropyl)-3-ethylcarbodiimide hydrochloride (EDAC) (116 μg, 60mM), and sulfo-N-hydroxysuccinimide (sulfo-NHS) (132 μg, 60 mM) in a volume of 10 μL DMF was rotated for 3 hours at 25 °C in the dark, in order to activate the three carboxyl groups of C*e*6. The amount of C*e*6 used corresponded to a 1000-fold molar excess of this agent relative to the available amino groups present in 800 μg of GVs. Separately, 50 μL of DMF was added to 800 μg of GVs suspended in 60 μL of PBS at pH 7. The activated C*e*6 solution (10 μL) was then added to the GV suspension and the final mixture was rotated for 5 hours in the dark at 25 °C. The nanoparticles were subsequently separated by centrifugation (300 g, 25 °C) and washed with PBS to remove residual free C*e*6. The grey-colored, C*e*6-modified GV suspension was stored at 4 °C in PBS.

### Determination of C*e*6 loading on GVs

The amino acid composition and protein concentration of the C*e*6-GV preparation were accurately determined by amino acid hydrolysis (Hospital For Sick Children Toronto, CA). C*e*6-GV samples were also hydrolyzed (110 °C in 6N HCl *in vacuo)* to release C*e*6 covalently bound to lysine ε-amino groups on the surface of GVs. The concentration of C*e*6 released from C*e*6-GV sample hydrolysates was determined by fluorescence spectroscopy using a free C*e*6 standard curve (λ_exc_ 400 nm; λ_em_ 660 nm; see Supplementary Figure S4).

### Nanoparticle Characterization

The shape and size of intact C*e*6-GVs were established by transmission electron microscopy (Phillips/FEI Tecnai Hillsboro OR) and by dynamic light scattering (Zetasizer Nano ZS, Malvern, UK). The size distribution of C*e*6-GVs derived from electron micrographs was estimated using ImageJ software. ^28^ Emission spectra of C*e*6-GVs were recorded using a fluorescence microplate reader (Synergy H1) using a concentration of 1.9 μM of C*e*6 in the form of free C*e*6 or C*e*6-GV or the equivalent amount of WT GV. The Zeta potential of C*e*6-GV or WT GV nanoparticles (final concentration of 40 pM in 990μL of distilled water) was determined using the Zetasizer Nano ZS (Malvern, UK).

### Cellular uptake studies

For cellular studies, the human breast cancer cell line MCF-7 was purchased from ATCC (cat.# HTB-22). The human hypopharyngeal cancer cell line FaDu-GFP (AntiCancer Inc., San Diego, CA) was a gift from Dr. David Goertz (Sunnybrook Research Institute, Toronto). MCF-7 and FaDu-GFP cells (10^6^ cells) were seeded into wells of 12-well plates and incubated overnight (37 °C, 5% CO_2_) to enable cell attachment. C*e*6-GVs, free C*e*6 (both adjusted to 670 nM C*e*6) or native GVs (equivalent to the C*e*6-GV concentration) were dispensed into wells. Plates were then kept rotating at 4 °C or 37 °C in an incubator with cells being collected over time [0.5 to 48 hours] to assess their viability and the internalization of C*e*6 and GVs. Briefly, cells were trypsinized and washed with PBS once. Cellular uptake of C*e*6 as a function of incubation time was measured by flow cytometry as mean fluorescence intensities (MFI); ◻_exc_ 405 nm, ◻P Filter 660-2 nm, Dichroic 635LP; LSR II, BD Biosciences, San Jose, CA). Internalization events of C*e*6 and C*e*6-GVs in MCF-7 or FaDu-GFP cells were also recorded by confocal microscopy. Specifically, MCF-7 or FaDu-GFP cells were seeded into an 8-well confocal chamber at a density of 10^5^ cells for 24h (37 °C, 5 % CO_2_). The cells were then treated with C*e*6-GV, free C*e*6, or an equivalent amount of WT GV for 8 hours. Cell nuclei were stained using the cell permeable dye Hoescht 33342 (1 μM; ThermoFischer). Cells were then washed with PBS and both phase contrast and fluorescence (◻_exc_ 403 nm, ◻_em_ 663-738 nm) images captured using a Nikon A1 laser-scanning microscope.

### Cell viability measurements

The cytotoxicity of C*e*6-GV, free C*e*6, or WT GV towards MCF-7 human breast cancer and FaDu-GFP pharyngeal cancer cell lines was assessed using the tetrazolium salt-based WST-1 cell proliferation assay. Briefly, cells were seeded at an initial density of 10^5^ cells and were incubated overnight (37 °C, 5 % CO_2_) to enable attachment. The next day, cells were treated for 24 hours with serial dilutions of either C*e*6-GVs, equivalent molar doses of free C*e*6, or corresponding amounts of native GVs relative to C*e*6-GVs. Cells were subsequently washed once with PBS and exposed to a LED light source (660 nm) for 10 min (ABI 25W Deep Red). The source irradiance was determined to be 15-35 mW/cm^2^ (Newport Powermeter 1918-R). The plate containing C*e*6-GV, free C*e*6, or WT GV-treated cells not exposed to the light source served as a control for non-light associated cytotoxicities. Following light exposure, cells were incubated for an additional 24 hours in the dark. After this period, media was discarded and cells were incubated with 10 μL of WST-1 reagent (Roche) and 90 μL of growth medium for 4 hours. Absorbance readings at 480 nm were then recorded using a microplate reader (Synergy H1). Cell viability was normalized relative to a positive control (50 μg/mL Doxorubicin-HCl treatment leading to complete cell death) and a negative control (untreated cells) using the following equation:

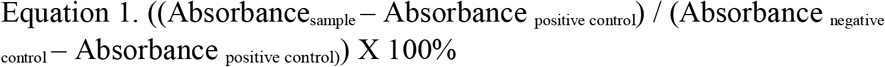

Two independent trials were performed and each data point represents the average cell survival percentage (± SD) derived from proliferation assays performed in triplicate for each tested treatment.

### Detection of intracellular reactive oxygen species (ROS)

2,7-Dichlorofluorescein diacetate (DCF-DA) (Sigma) is a non-fluorescent, cell-permeable probe used to measure intracellular ROS. DCF-DA is deacetylated by intracellular enzymes to a non-fluorescent dye that is finally oxidized by ROS to the fluorescent compound dichlorofluorescein. ^29^ For this assay, MCF-7 or FaDu-GFP cells were seeded into 24-well plates (5 × 10^5^ cells) overnight and were subsequently treated for 8 hours with C*e*6-GV, free C*e*6 (based on a molar concentration equivalent of 500 nM) or WT GV (at a dose identical to that of C*e*6-GV) to allow for their internalization. Cells were then washed with PBS and the plates were either exposed to light (660 nm) for 10 min or kept in the dark. After the light treatment, 2.5 μl of 1 mM DCF-DA dissolved in DMSO (5 μM final concentration in wells) was dispensed into each well and left for 30 min at 37 °C in an incubator. Cells were washed twice under dark conditions with PBS and the DCF fluorescence emission signal was recorded using a micro-plate reader (λ_exc_ 488 nm; λ_em_ 525 nm) (Synergy H1). The fluorescence emission values were normalized relative to the average DCF fluorescence of cells treated with free C*e*6 in the absence of light. The results were analyzed with a one-tailed paired t-test to evaluate the difference between paired groups based on sample size of n =10. Significant differences were determined as α < 0.05. Normality of the data distribution was confirmed using Kolmogorov-Smirnov test.

## Data Availability

Data is available upon reasonable request from the corresponding author.

## Acknowledgements

We would like to thank Dr. Gang Zheng (University of Toronto) for critically reviewing this manuscript and Dr. Christine Allen (University of Toronto) for her support with this project. This study was supported by a grant from the Canadian Institutes of Health Research to J.G.

## Author Contributions

A.F designed the experiments with advice from J.G. A.F. performed the experiments, analyzed the results, and prepared the manuscript. The manuscript was revised by J.G.

