## Supporting Information for "Light-Activated Nanoscale Gas Vesicles Selectively Kill Tumor Cells"

### Methods

#### Mass Spectrometry Analysis of Native GVs

LC-MS/MS was locally performed on purified wild type GVs at the SPARC Molecular Analysis Facility (Sick Kids, Toronto, ON, CAN)

#### Cytotoxicity of Native GVs

The cytotoxicity of native GVs towards MCF-7 and FaDu-GFP cell lines was calculated from cell viability measurements performed as described in the Material and Methods Section.

#### Construction of Standard Curve of Fluorescence signal as a function of Free C*e*6 Concentration

C*e*6 was dissolved in ethanol and fluorescence signals ($\lambda_{exc}$ 400 nm; $\lambda_{em}$ 660 nm) were recorded (in triplicate) as a function of free C*e*6 concentrations using a Synergy H1 microplate reader.

### Results

#### Mass Spectrometry Analysis of Native GVs

LC-MS/MS analyses were performed on purified native *Halobacterium* GVs,to identify the most abundant protein species. Figure S1 provides a list of the most common protein species observed, highlighting GvpA as the main component of GVs. The only other GV protein detected was the non-structural protein GvpD. No GvpC peptide was observed by mass spectrometry suggesting that this subunit was removed during purification steps. The normalized total spectra were determined by multiplying the spectral count for individual proteins by the ratio of the average total spectra for all samples to the number of spectra for one particular sample.

| Name of Protein | Normalized Total Spectra (%) |
|---|---|
| Gas Vesicle Protein A [Halobacterium salinarum NRC-1] | 28 |
| H+-transporting ATP synthase subunit K [Halobacterium salinarum NRC-1] | 7 |
| Thermosome subunit beta [Halobacterium salinarum NRC-1] | 5 |
| Conserved hypothetical protein [Halobacterium salinarum NRC-1] | 4 |
| Cell surface glycoprotein [Halobacterium salinarum NRC-1] | 4 |
| Bacteriorhodopsin [Halobacterium salinarum NRC-1] | 3 |
| Conserved hypothetical protein [Halobacterium salinarum NRC-1] | 3 |
| Hypothetical protein VNG_1802H [Halobacterium salinarum NRC-1] | 3 |
| Dipeptide ABC transporter dipeptide-binding [Halobacterium salinarum NRC-1] | 2 |
| RepJ (plasmid) [Halobacterium salinarum NRC-1] | 2 |

**Supplementary Figure S1**. LC-MS/MS was performed on purified wild type GVs and identified GvpA as the main component in the preparation. The normalized total spectra is calculated by multiplying the spectral count for each protein by the ratio of the average total spectra for all samples to the number of spectra for one particular sample.

Figure S2 shows the calculation used to estimate the molecular weight of a single *Halobacterium* GV.

<u>Estimating the surface area (SA) of one WT-GV</u>:
<u>Assumption:</u> The structure of a wild type (WT) GV approximates the shape of a Prolate Spheroid with semi-axes of **a** =129 nm (257/2) and **b** =190 nm (379/2) based on TEM results (Table 1). The <u>surface area</u> of one WT-GV was calculated to be 277500 $nm^2$ according to the following equation:

$$SA = 2\Pi\ (a^2+[(a \times b \times e) / \sin(e)])$$

Where e = arc cos(a / b)

The <u>surface area</u> of a single GvpA subunit ($SA_{GvpA}$) was reported to be 4.6 X 1.1 $nm^2$ [≈ 5 $nm^2$] (Blaurock and Walsby 1976). Using these values, the approximate number of GvpA molecules per GV was estimated to be 55,500 [$SA_{GV}$ / $SA_{GvpA}$ = 277 500 $nm^2$ / 5 $nm^2$].

Since WT GVs are mainly composed of GvpA subunits (>90% of the nanobubble core), the projected mass of a single GV is ~ 55,500 GvpA subunits x 8.01 kD/GypA or 444 MDa.

**Supplementary Figure S2.** Estimating the number of GvpA proteins per GV and the molecular weight of a single GV.

**Cytotoxicity of Native GVs**

The toxicity of native GVs towards MCF-7 and FaDu-GFP cancer cell lines in the presence and absence of red light was evaluated using the WST-1 cell proliferation assay. As shown in Figure S3, the results demonstrated a lack of toxicity towards GVs at most doses tested.

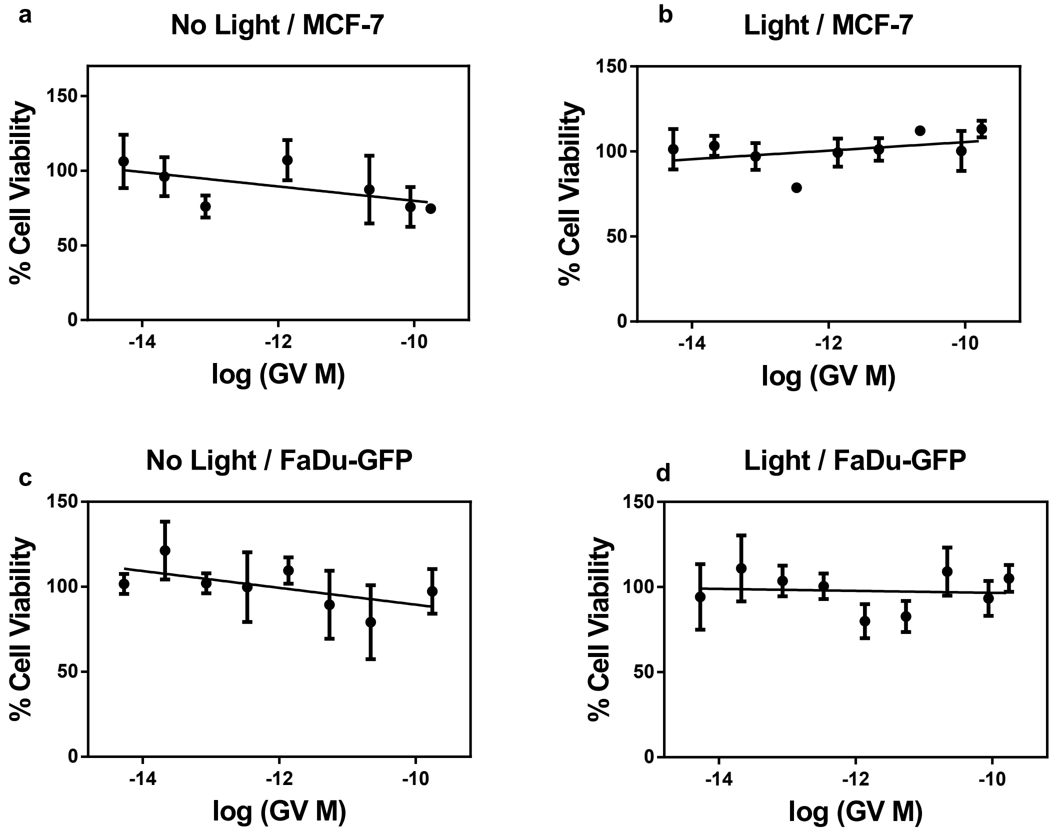


**Supplementary Figure S3.** Toxicity of MCF-7 or FaDu-GFP cells towards Native GVs as determined by WST-1 viability assay. The GV concentration range (X axis) reflects the molar concentration of GVs relative to C*e*6 based on the estimate that ~60,000 C*e*6 molecules are coupled to each wild type GV (Fig. 6)

**Construction of Standard Curve of Free C*e*6 Fluorescence versus Concentration**

A calibration curve of Free C*e*6 fluorescence as a function of concentration was derived to quantify the amount of C*e*6 loaded on GVs as described in the manuscript.

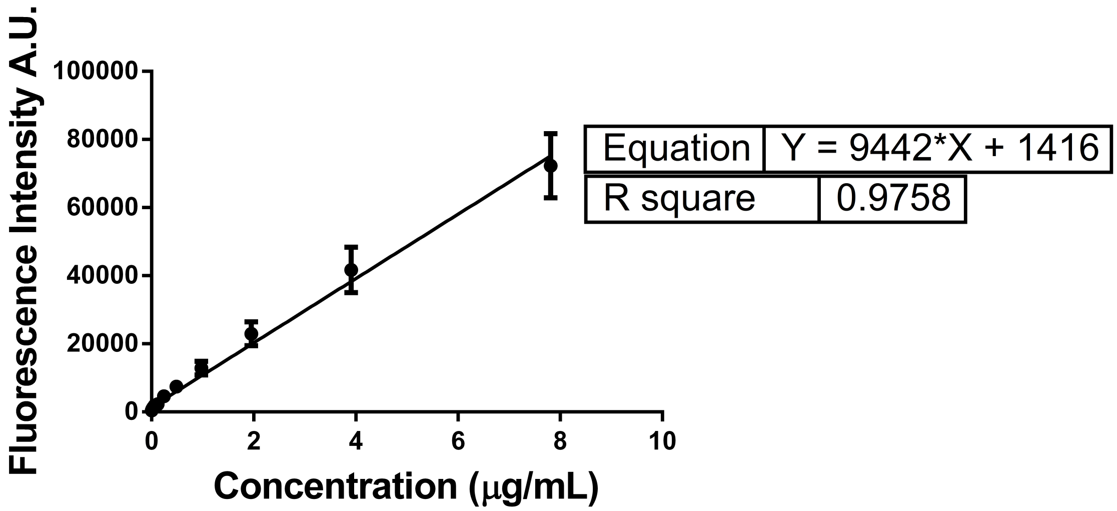


**Supplementary Figure S4**. Free C*e*6 fluorescence intensity as a function of concentration.
